# A high-throughput assay to identify robust inhibitors of dynamin GTPase activity

**DOI:** 10.1101/153106

**Authors:** Aparna Mohanakrishnan, Triet Vincent M. Tran, Meera Kumar, Hong Chen, Bruce A. Posner, Sandra L. Schmid

## Abstract

Clathrin-mediated endocytosis is the major pathway by which cells internalize materials from the external environment. Dynamin, a large multidomain GTPase, is a key regulator of clathrin-mediated endocytosis. It assembles at the necks of invaginated clathrin-coated pits and, through GTP hydrolysis, catalyzes scission and release of clathrin-coated vesicles from the plasma membrane. Several small molecule inhibitors of dynamin’s GTPase activity, such as Dynasore and Dyngo-4a, are currently available, although their specificity has been brought into question. Previous screens for these inhibitors measured dynamin’s stimulated GTPase activity due to lack of sufficient sensitivity, hence the mechanisms by which they inhibit dynamin are uncertain. We report a highly sensitive fluorescence-based assay capable of detecting dynamin’s basal GTPase activity under conditions compatible with high throughput screening. Utilizing this optimized assay, we conducted a pilot screen of 8000 compounds and identified several “hits” that inhibit the basal GTPase activity of dynamin-1. Subsequent dose-response curves were used to validate the activity of these compounds. Interestingly, we found neither Dynasore nor Dyngo-4a inhibited dynamin’s basal GTPase activity, although both inhibit assembly-stimulated GTPase activity. This assay provides the basis for a more extensive search for robust dynamin inhibitors.

## Introduction

Dynamin is a large multidomain GTPase known for its role in catalyzing membrane fission in clathrin-mediated endocytosis (CME) [1-3]. It consists of five functional domains: the N-terminal GTPase domain (G domain); the middle domain and the GTPase effector domains (GEDs), which together form the stalk of dynamin; a pleckstrin homology (PH) domain; and the C-terminal proline- and arginine-rich domain (PRD), which interacts with many SH3 domain-containing proteins [4]. Dynamin assembles at the necks of invaginated clathrin-coated pits and catalyzes scission and release of clathrin-coated vesicles from the plasma membrane. Dynamin is recruited to nascent coated pits in its unassembled state and also plays a regulatory role during the early stages of CME [5-7].

Most GTPase family members that function as regulatory proteins do so by switching between GTP-bound ‘active’ conformations and GDP-bound ‘inactive’ states. Their intrinsic GTP hydrolysis rates are slow, and rate-limited by the exchange of tightly-bound GDP for GTP. These two steps in the GTP hydrolytic cycle are regulated by GTPase activating proteins (GAPs) and GTP exchange factors (GEFs), respectively. In this regard, dynamin is an atypical GTPase as it has a low affinity for GTP (2-5 μM), a high rate of GDP dissociation (~ 60-90 s^-1^), and a comparatively robust and measurable basal rate of GTP hydrolysis (~ 1 min^-1^ at 37°C) [8]. However, upon self-assembly, interactions between G domains can stimulate GTPase activity *in trans* [9]. *In vivo*, dynamin self-assembles into short helical structures that surround the necks of deeply invaginated coated pits. *In vitro,* dynamin assembles into long helical arrays around lipid nanotubes whereby its GTPase activity is stimulated > 100-fold [10]. Dynamin’s GTPase activity can also be stimulated, albeit to a lesser extent, through interactions with divalent SH3 domain containing partners such as Grb2 [11,12] or under low salt conditions that favor dynamin self-assembly [13].

Given its importance for clathrin-mediated endocytosis, coupled to the fact that it is one of the few enzymes known to be required for CME, small molecule inhibitors of dynamin’s GTPase activity have been sought as potentially powerful tools for studying CME. Indeed, several chemical inhibitors of dynamin have been reported and are commercially available, including Dynasore [14,15] and its structural derivative, Dyngo-4a [14,15]. However, recent findings have brought into question the specificity of these compounds. For example, Dynasore and Dyngo-4a continue to inhibit endocytosis in triple dynamin-1, 2, and 3 knockout cells, thus revealing potential off-target effects [16]. These off-target effects on endocytosis may reflect their reported ability to perturb plasma membrane cholesterol levels [17] and destabilize actin filaments [16]. Recently, Dynasore was shown to impair VEGFR2 signaling in an endocytosis-independent manner [18]. Based on the clear evidence for dynamin-independent, off-target effects of these compounds, there remains a need to develop more specific and robust dynamin inhibitors.

Previous screens for small molecule inhibitors of dynamin’s GTPase activity were based on the detection of released phosphate using a malachite green colorimetric assay. However, this assay lacks sufficient sensitivity to detect dynamin’s basal GTPase activity, especially when measured at room temperature and at the low concentrations of dynamin and GTP practically needed for the design of a high-throughput assay. To circumvent this, previous high throughput screens measured dynamin’s stimulated GTPase activity either in the presence of GST-Grb2 [14,15] or with sonicated phosphatidylserine (PS) liposomes at low salt [14,15]. Dynasore, and by extension Dyngo-4a, was shown to be a noncompetitive inhibitor of dynamin’s GTPase activity [14,15]. Hence its mechanism of action, which remains unknown, may reflect indirect effects on dynamin assembly or aggregation.

Here we report the optimization of a new, highly sensitive, and robust HTS-compatible assay to detect the basal GTPase activity of dynamin and its validation in a preliminary screen of 8000 compounds.

## Materials and methods

### Dynamin expression, purification, and preparation

Dynamin-1 (Dyn1) was expressed in Sf9 insect cells and purified by affinity chromatography, as previously described [19]. Protein aliquots were flash frozen in liquid nitrogen and stored in -80°C in 5% v/v glycerol. Prior to running assays, frozen aliquots of Dyn1 were thawed and centrifuged at 100,000 g for 15 minutes to remove any aggregates. Dyn1 concentration was determined by measuring its absorbance at 280 nm with a UV/Vis spectrophotometer (Beckman Coulter Inc.) using a molar absorptivity coefficient of 73,800 M^-1^ cm^-1^.

### Transcreener GDP fluorescence polarization assay

The Transcreener^®^ GDP FP (BellBrook Labs) assay is an immune-competition assay based on a mouse monoclonal antibody that selectively binds Alexa633-conjugated GDP over GTP. The preformed Alexa633-GDP antibody complex is added at the end of the reaction, and the GDP generated displaces bound Alexa633 fluorescent tracer, resulting in a decrease in its far-red fluorescence polarization (FP). The assay detects GDP production with high sensitivity at low substrate concentrations and has been used to develop an HTS compatible assay for the enzymatic activity of ARFGAP [20]. Stocks of 5 mM GTP, 10x Stop & Detect Buffer B, 400 nM GDP Alexa633 Tracer, and 3100 μg/mL GDP antibody were purchased from BellBrook Labs (Madison, WI). Assays were performed in Corning 384-well plates at room temperature.

All assays were conducted in the endpoint format, which measures total GDP production after 60 minutes of incubation at room temperature. The optimized reaction (total volume 15 μL) was initiated by adding 5 μL of GTP (3x stock of 30 μM, final concentration 10 μM) to wells containing 10 μL Dyn1 (1.5x stock, final concentration 50 or 100 nM). Reactions were terminated after 60 minutes with the addition of 5 μL of GDP detection mixture (4x stock of 8 nM GDP Alexa633 Tracer, 40 mM HEPES, 80 mM EDTA, 0.04% Brij and 34.4 μg/mL GDP antibody). The plates were incubated for another 60 minutes, allowing the GDP antibody-GDP binding to reach equilibrium. The plate was then read for fluorescence polarization in millipolarization units (mP) using an EnVision multi-modal microplate reader (PerkinElmer, Inc.). The mP values of the reaction-containing wells were subtracted from those containing no enzyme to obtain ΔmP values. To convert the polarization data to GDP released, a standard curve representing 0 to 100% GDP conversion from 10 μM GTP was generated. Using GraphPad Prism, the ΔmP values were fitted to the standard curve to obtain the total GDP production. Importantly, the assay was insensitive to DMSO concentrations up to 3%.

### 8000-compound library screen

A screen was performed using the optimized Transcreener GDP FP assay on a diverse library subset of 8000 small molecule compounds, provided by the UT Southwestern HTS Core. For the HTS screen, the final concentration of Dyn1 in the reaction mixture was 50 nM. 0.3 μL of compound in 100% DMSO (final concentration 10 μM compound, 2% DMSO) was added to the Dyn1 and pre-incubated for 30 minutes. Controls for the screen included reactions that lacked enzyme (positive control for inhibition) and uninhibited reactions containing only DMSO (negative control for inhibition), which were dispensed in single columns in each plate. Solutions were dispensed using automated liquid handling devices.

### Data analysis

The primary screen data were analyzed using Genedata Screener^®^ software. The Z’ factors for the mock screen and the 8000-compound pilot screen were calculated using the equation below:
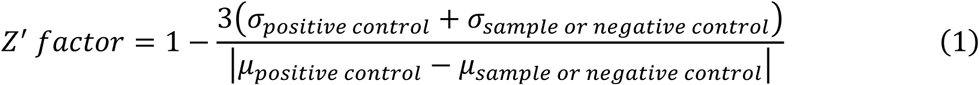

where σ_positive control_ is the standard deviation of the positive controls for inhibition, and σ_sample or negative control_ is the standard deviation of the samples or negative controls for inhibition, respectively. μ_positive control_ is the mean of the positive control for inhibition, and μ_sample or negative control_ is the mean of the samples or neutral DMSO controls, respectively.

The samples were normalized by a two-point correction method using the equation below:
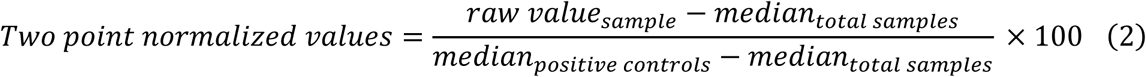

where median_total samples_ is defined as the median of all library compound-containing reaction wells within the plate.

The two-point normalized activity values were adjusted using a correction factor to account for systematic errors within and across assay plates [21]. The correction factor of a well in a given plate is calculated using pattern detection algorithms that are proprietary to the Screener^®^ software (Genedata, Inc.). The corrected activity values were then used to determine the robust Z (RZ) score with the following equation:
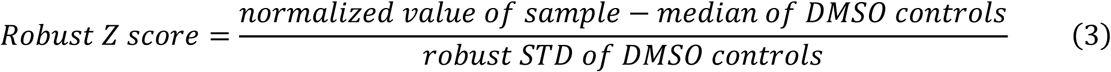

where robust STD is the standard deviation calculated using the median of the DMSO controls (negative control for inhibition).

For the confirmation screen and dose response curves, the data were analyzed by normalizing the sample GDP released to the control GDP released using the following equation:
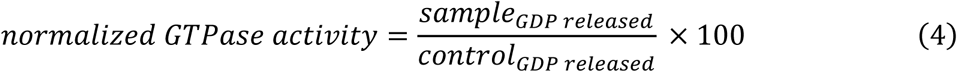

### Malachite green assay

The lipid nanotube (NT)-stimulated malachite green assays were performed in 96-well plates at 37°C. The final reaction consisted of 100 nM Dyn1, 25 μM GTP, and 300 μM lipid nanotubes. The assay and reagent preparations were performed according to our published protocol [22]. All general chemicals were purchased from Sigma-Aldrich (St. Louis, MO). Compounds tested in the malachite green assay were purchased from Chembridge and ChemDiv (both located in San Diego, CA).

## Results

### Optimization of a fluorescence polarization assay to detect basal GTPase activity of dynamin

Fluorescence polarization (FP) is a method that allows for rapid and quantitative analysis of diverse molecular interactions and enzyme activities [23]. Polarization measures the change in the molecular movement of the labeled species. It is the ratio of the difference between the vertical and horizontal components of the emitted light over their sum [20]. In recent years, FP has been successfully used in HTS of compound libraries to identify small molecule inhibitors of protein-protein interactions.

Bellbrook labs has developed an assay that detects GDP using a competitive FP immunoassay. GDP released upon hydrolysis of GTP by GTPases displaces a fluorescent tracer from the antibody, resulting in a decrease in polarization due to increased rotational mobility. The antibody has 140-fold specificity for GDP versus GTP, which allows sensitive measurement of GDP in the presence of excess GTP. Given that the antibody used has a finite selectivity for GDP over GTP, it was necessary to determine the optimal concentration of the Alexa-GDP antibody conjugate needed for maximum mP measured in the presence of 10, 100 and 500 μM GTP, our initial substrate concentrations. For this purpose, the Alexa-GDP antibody conjugate was titrated into the reaction mixture containing GTP (10, 100 or 500 μM) and assay buffer (20 mM HEPES, 150 mM KCl, 1 mM EDTA, 1 mM EGTA and 1 mM DTT, pH 7.4). The data were fitted to a variable slope sigmoidal dose-response curve using GraphPad Prism (Fig 1A). From the titration curves, we determined the optimal GDP antibody concentrations to be 8.6, 81.5 and 405.5 μg/mL for 10, 100 and 500 μM GTP, respectively (highlighted data points in Fig 1A). These concentrations were chosen near signal saturation and represent a good compromise between sensitivity and maximal polarization value.

**Fig 1.**
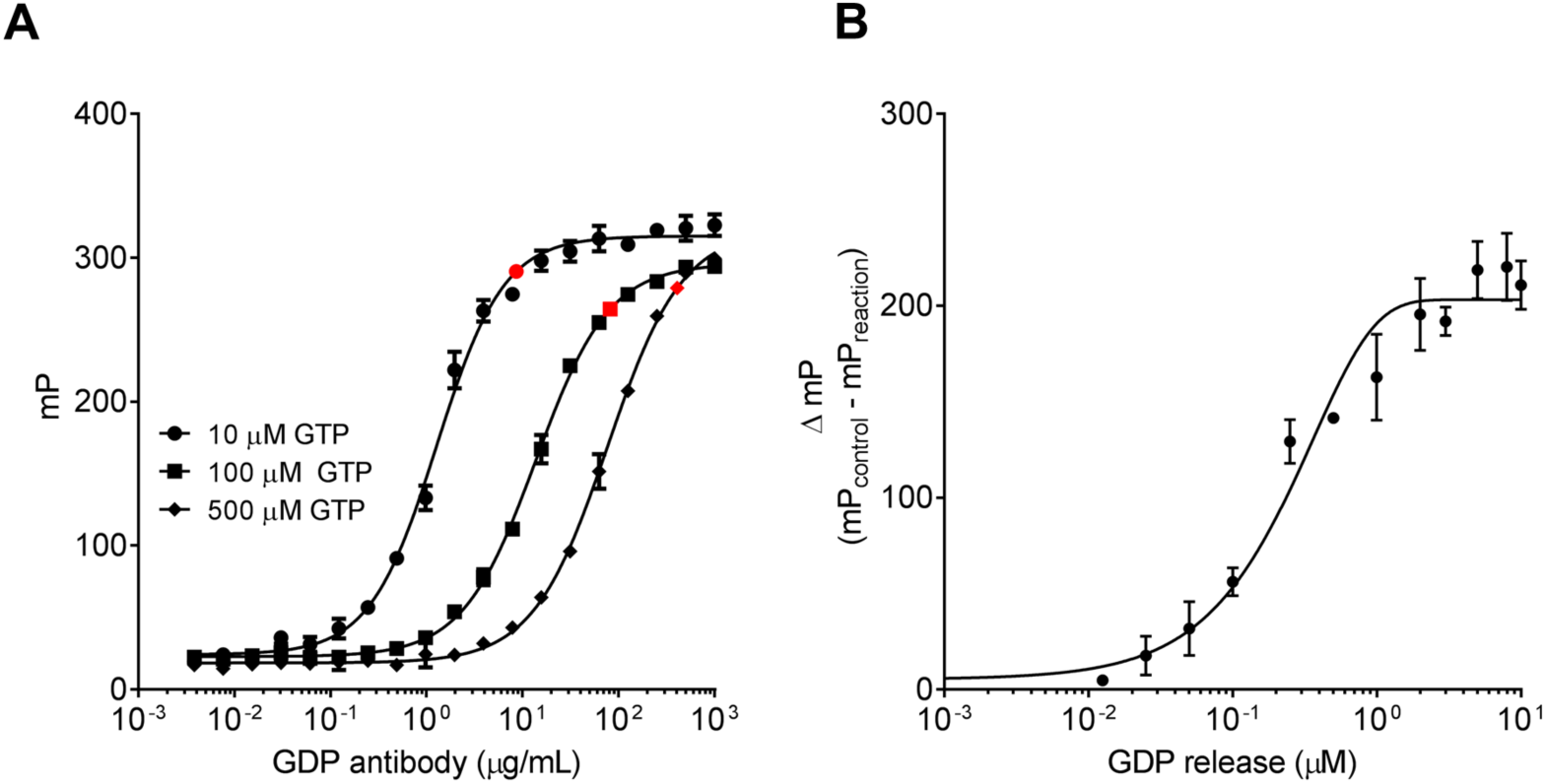
Optimization of assay sensitivity to measure GDP production. **A)** The GDP antibody was titrated to determine its optimal concentrations for 10, 100 and 500 μM GTP. Optimal antibody concentrations are represented by the highlighted points (n = 1, measured in triplicates). **B)** Standard curve representing the conversion of 0 to 100% GDP from 10 μM GTP. This curve was used to convert the fluorescence polarization data to GDP released (n = 1, measured in triplicates). Data are presented as mean ± SD.

To convert ΔmP to μM GDP released, we generated a standard curve by titrating increasing concentrations of GDP in the presence of GTP to mimic reaction conditions. The assay accurately measures GTP hydrolysis in the range of 0.05% to 10% of the substrate converted (Fig 1B).

To determine the optimal conditions for high throughput screening, we measured ΔmP for increasing concentrations of Dyn1 (0.3 nM to 5000 nM) at three different concentrations of GTP (10, 100, and 500 μM) (Fig 2A). These titrations established that 50 nM Dyn1, assayed in the presence of 10 μM GTP for 60 minutes, resulted in excellent signal-to-noise with high reproducibility. We further confirmed that, under these conditions, the basal rate of GTP hydrolysis by Dyn1 (~ 0.04 min^-1^ at 10 μM GTP) was linear for 60 min (Fig. 2B). These results are consistent with assays performed at room temperature and under low substrate concentrations. We chose 60 minutes to ensure that substrate consumption remained below 10%. Importantly, no signal was detected at any concentration of GTP when S45N mutant Dyn1, which is unable to bind GTP [24], was used as a negative control (Fig 2C). We further confirmed that under these conditions, GTPase activity is directly proportional to the concentration of Dyn1. Thus, there is no evidence of cooperativity and the assay measures the basal rate of GTP hydrolysis of unassembled Dyn1 (Fig 2D).

**Fig 2.**
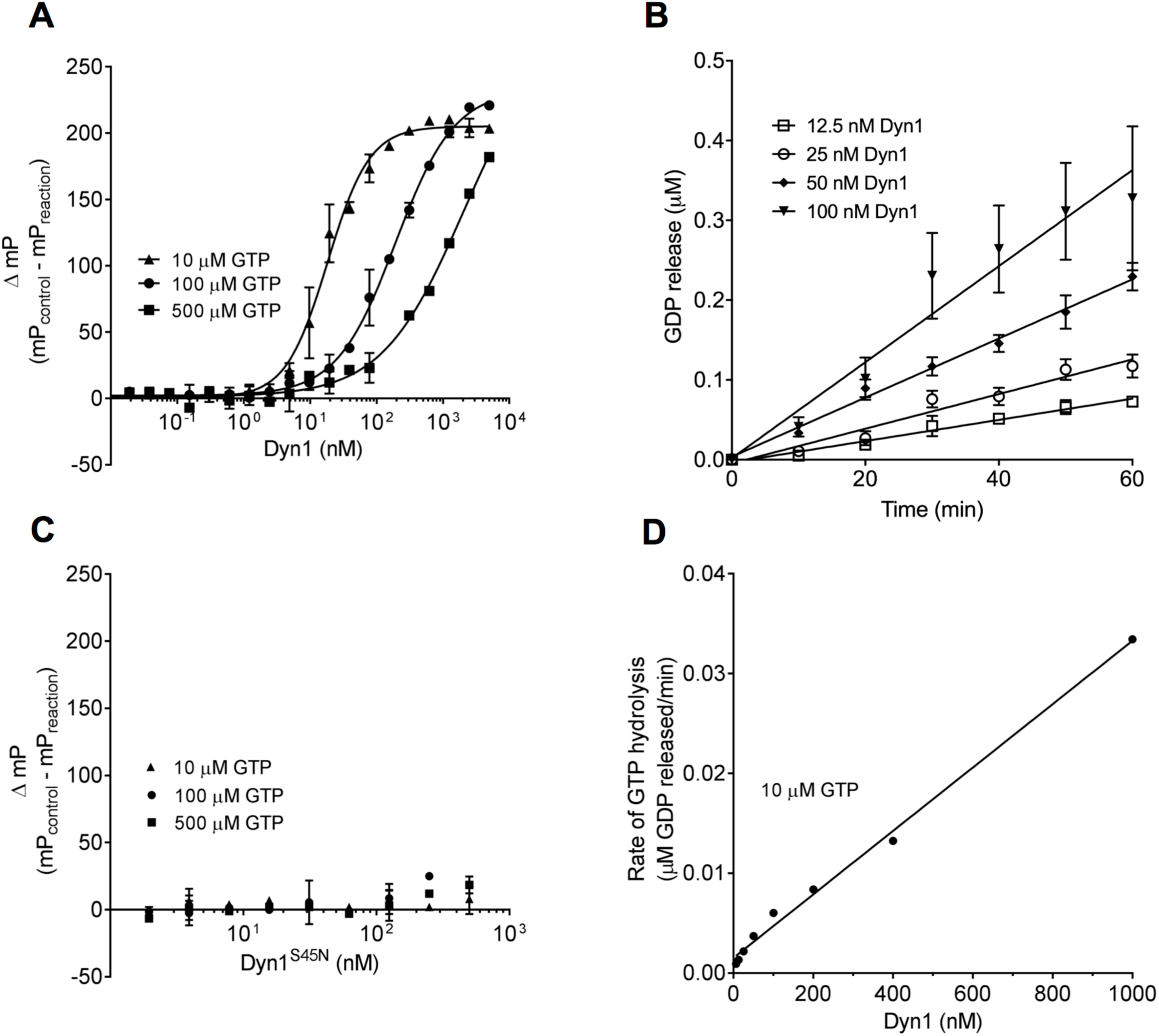
Detection of basal GTP hydrolysis by Dyn1. **A)** Dyn1 was titrated from 0.3 nM to 5000 nM in the presence of 10 μM, 100 μM or 500 μM GTP and GTP hydrolysis was measured as ΔmP after 60 min incubation at room temperature (n = 1, measured in triplicates). **B)** The GTP hydrolysis by Dyn1 was linear during 60 minute incubations at room temperature (n = 4, each measured in duplicates). **C)** The GTP binding mutant, Dyn1^S45N^, shows no activity at any concentration of GTP (n = 1, measured in triplicates). **D)** The basal GTPase activity is proportional to Dyn1 concentration from 0 to 1000 nM, indicating no cooperativity under these concentrations (n = 3, each measured in triplicates). Data are presented as mean ± SD.

### Pilot screen, hit selection and validation

We measured the robustness of the assay under our optimized HTS conditions to determine whether ‘hits’ could be identified with high confidence. mP values obtained from 15 μL reactions after a 60-minute incubation with 50 nM Dyn1 and 0.3 μL of 100% DMSO in the presence of 10 μM GTP were compared to those lacking Dyn1 (Fig 3A). The average Z’ factor for the mock screen, which was calculated to be 0.56, indicated that the assay was sufficiently robust for screening purposes.

**Fig 3.**
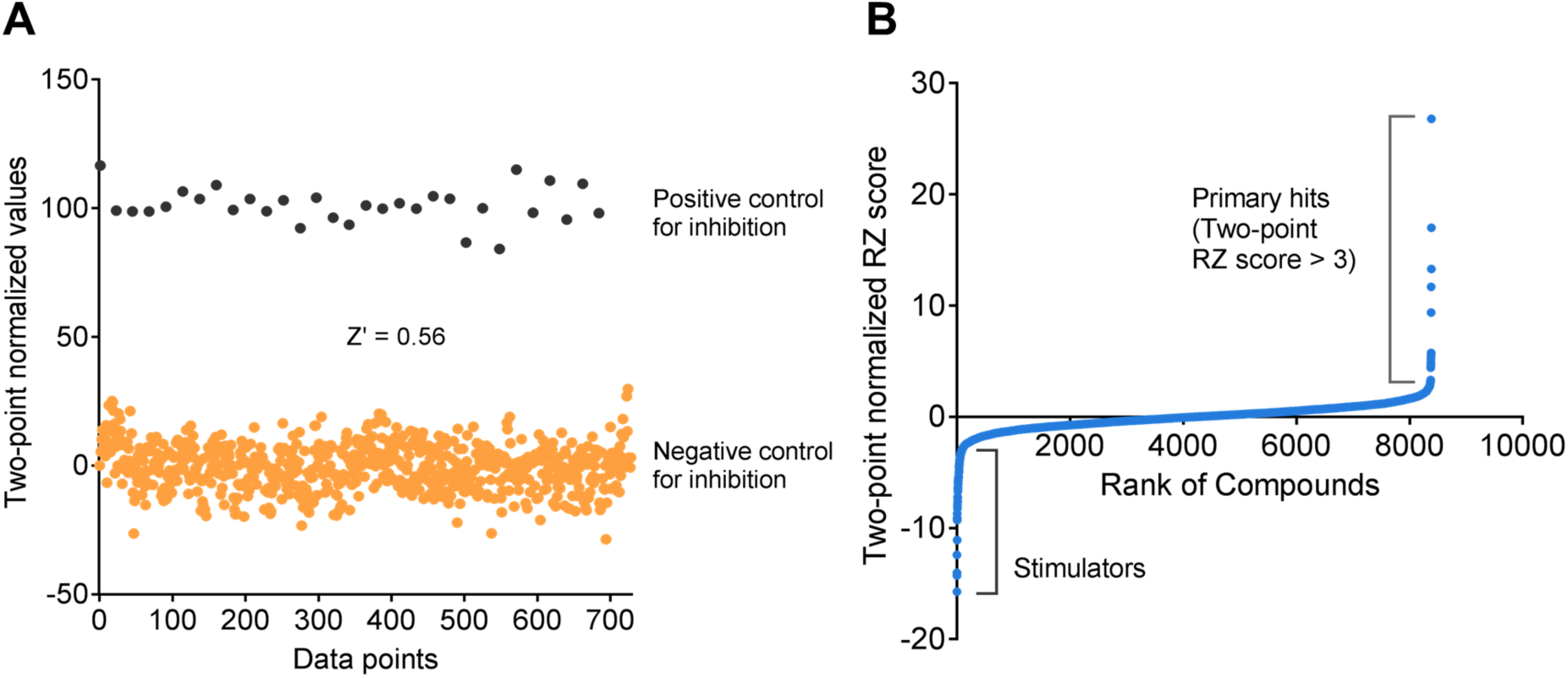
Assay validation and application for high-throughput screening of 8000 compounds. **A)** The assay was validated in a preliminary mock screen that compared reactions containing no enzyme (positive control for inhibition) to uninhibited reactions containing DMSO (negative control for inhibition). **B)** A quantile-quantile (Q-Q) plot of the pilot screen of 8000 compounds. The compounds are ranked from 0 to 8000 according to their two-point robust Z score. Primary hits were chosen based on a RZ score of greater than 3.

A pilot screen using an 8000-compound diversity subset of the chemical library at UT Southwestern was conducted using the optimized Transcreener GDP FP assay. The compounds were tested for their inhibitory effects on the Dyn1 GTPase activity at a concentration of 10 μM.

Intrinsically fluorescent compounds, which alter total fluorescence intensity per well, were eliminated, as they would impact analysis. After careful analysis of the data, we identified 42 compounds with a robust Z score greater than 3 as primary hits (Fig 3B).

These compounds were re-tested in a confirmation screen at three different concentrations to validate their inhibitory effects, yielding 4 promising compounds based on concentration-dependent inhibition. To confirm the inhibitory effects of the 4 compounds, we conducted 11-point dose response curves, with concentrations ranging from 1 nM to 100 μM (Fig 4A). The IC50 values of these compounds ranged from < 1 μM to > 50 μM. We focused on compound 24 which had an IC_50_ of 0.58 μM.

**Fig 4.**
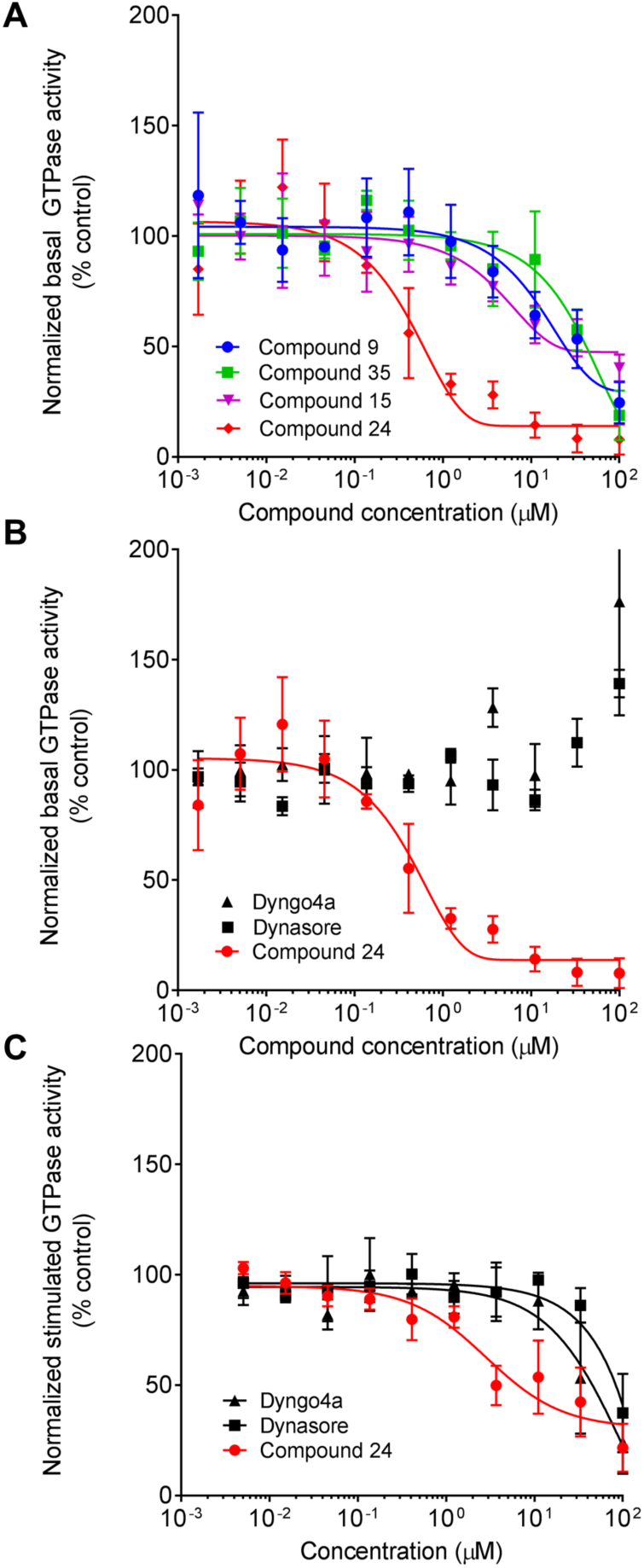
Comparison of commercially available inhibitors with compound 24. **A)** 11-point dose response curve of the 4 validated hits from the primary screen of 8000 compounds measured using 50 nM Dyn1 and 10 μM GTP (n = 1, measured in triplicates). **B)** Dynasore and Dyngo-4a were tested alongside compound 24 in the Transcreener assay (n = 1, measured in triplicates). **C)** Dose response curves for the inhibitory effects of Dynasore, Dyngo-4a and compound 24 on the lipid nanotubestimulated GTPase activity of 100 nM Dyn1 assayed in the presence of 300 μM lipid nanotubes and 25 μM GTP and measured using the malachite green assay (n = 3, each measured in triplicates). Data are presented as mean ± SD.

### Secondary assay and comparison with Dynasore and Dyngo-4a

Compound 24 was compared with the two commercially available dynamin inhibitors, Dynasore and Dyngo-4a, in a dose-response assay measuring inhibition of basal GTPase activity under high salt (150 mM KCl) conditions. The concentration of inhibitors ranged from 1 nM to 100 μM. As seen in Fig 4B, Dynasore and Dyngo-4a do not appear to inhibit basal GTPase activity even at high concentrations, in contrast to previous findings in which assays were performed under conditions that measure dynamin’s stimulated, assembly-dependent GTPase activity [14,15]. Therefore, to more closely parallel previous studies we tested both commercial inhibitors in comparison to compound 24 for their effects on dynamin’s stimulated GTPase activity. Assays were performed in the presence of PI(4,5)P2-containing lipid nanotubes (NT), whose diameter (~ 20 nm) resembles the neck of an invaginated coated pit [10], using the malachite green assay. Under these conditions (100 nM Dyn1, 300 μM lipid nanotubes, 25 μM GTP), both Dynasore and Dyngo-4a inhibited the NT-stimulated GTPase activity of Dyn1 with IC_50_ values of 83.5 and 45.4 μM, respectively, as compared to compound 24, which exhibited an IC_50_ of 6.4 μM in this assay (Fig 4C).

## Discussion

We have optimized a robust, high-throughput assay to measure the basal GTPase activity of unassembled Dyn1. This highly sensitive assay detects the release of low, nanomolar amounts of GDP and hence, accurately measures the intrinsic, basal rate of GTP hydrolysis, even at the low concentrations of dynamin and GTP necessary for HTS design and implementation. Previous high throughput screens using a less sensitive colorimetric malachite green assay to detect phosphate release were necessarily performed under conditions that stimulate dynamin’s GTPase activity, i.e. in the presence of dimeric GST-Grb2, which presumably aggregates dynamin, or with sonicated PS liposomes in low salt.

Utilizing the Transcreener GDP FP assay, we conducted an 8000-compound pilot screen and identified several compounds that inhibit the basal GTPase activity of Dyn1. The commercially available dynamin inhibitors, Dynasore and Dyngo-4a, were tested for their ability to inhibit dynamin’s basal GTPase activity in the Transcreener assay format. Although both Dynasore and Dyngo-4a could inhibit the NT-stimulated GTPase activity of Dyn1, neither was able to inhibit basal GTPase activity in our hands. Moreover, the reported IC_50_ values we measured for Dynasore and Dyngo-4a NT-stimulated GTPase activity performed at physiological salt concentrations using 100 nM dynamin were much higher than those reported for assays performed in the presence of sonicated PS liposomes, under low salt conditions with 20 nM dynamin (0.4 μM and 12 μM, respectively). Given the reported off-target effects of Dynasore and Dyngo-4a [16-18] and their uncertain mechanism of dynamin inhibition, a more robust and specific inhibitor of dynamin would be of immense value.

As with any assay, fluorescence polarization has its limitations. Compounds that are either auto-fluorescent, or affect the affinity of the anti-GDP antibody for the tracer may be misinterpreted as potential hits [20]. The hits must therefore be validated in secondary assays such as the malachite green assay and eventually for their ability to inhibit dynamin-dependent, clathrin-mediated endocytosis in intact cells.

Having validated our assay using an 8000-compound pilot screen, we are currently expanding our search for robust, specific, and cell-permeable dynamin inhibitors by screening the entire UT Southwestern chemical library of 230,000 compounds using the optimized Transcreener GDP FP assay.

## Acknowledgements

We thank Dana K. Reed for helping with protein expression and purification. We also thank the members of the Schmid lab for thoughtful discussions.

